# Assessing trait driver theory along abiotic gradients in tropical plant communities

**DOI:** 10.1101/2020.02.15.950139

**Authors:** Jehová Lourenço, Erica A. Newman, Camilla Rozindo Dias Milanez, Luciana Dias Thomaz, Brian J. Enquist

## Abstract

1. Despite the many studies using trait-based approaches to assess the impact of environmental gradients in forest trait composition, the relative roles of (i) intraspecific variation in community assembly and (ii) microclimatic or fine scale abiotic variation in shaping local trait diversity remain poorly understood. To advance their understanding we tested several assumptions and predictions of trait driver theory (TDT). We quantified the shape of trait distributions related to tree carbon, nutrient economics and stem hydraulics across a small-scale but steep gradient of soil water availability.
2. We utilized a unique and steep environmental gradient in the coastal Brazilian Atlantic forest (*restinga*) communities that spans a very short distance (207 ±60 meters). We collected leaf and wood samples of tree species across 42 patches (or plots) of *restinga* forest. Furthermore, to detect if species directionally shift in niche space, we analyzed species composition in multidimensional hypervolume space.
3. Despite short geographic distances, we observed large shifts in species replacement and intraspecific variation reflected by a directional shift in plant function. Consistent with TDT, we observe (i) trait distributions that are skewed in directions consistent with a forest responding to recent hotter and drier; (ii) peaked trait distributions, indicating strong functional convergence; and (iii) conditions decreasing means and variances of several leaf carbon and nutrient economic traits as well as stem hydraulic traits.
4. *Synthesis.* Observed species replacements along the water table gradient and interspecific measures of functional diversity (community kurtosis and skewness) are consistent with strong phenotype/environmental matching of plant carbon, nutrient, and hydraulic strategies. We observe environmental filtering in both extremities of the gradient, selecting for acquisitive (wet) to conservative (dry) setup of traits. Similarly, species that span the entire water availability gradient are characterized by directional intraspecific shifts in multi-trait space that mirror interspecific shifts. Strong environmental gradients across short spatial scales provide unique systems to accurately assess assembly processes and address long-held assumptions and timely hypothesis predicted by trait driver theory.

## Introduction

A central challenge in ecology is understanding and predicting the response of communities to changing climates, past, present, and future. The use of natural gradients to understand how biological diversity responses to climate has a long history (Clements, Hanson, & Weaver, 1929; Gause, 1932; Hutchinson, 1957). Observed ecological shifts along environmental gradients (such as temperature, water availability, and so on) can reflect evolutionary adaptation, plastic shifts in phenotypes, and responses to multiple stressors over a range of conditions (Polechová & Barton, 2015; Riesch, Plath, & Bierbach, 2018). Nevertheless, disentangling and deducing the mechanisms that drive shifts in biological diversity remains a fundamental problem (Adler, HilleRislambers, & Levine, 2007; Hubbell, 2001; McGill, 2003).

In this paper, we assess two central concepts in trait-based ecology. First, we utilize recent theoretical advances to assess the importance of phenotype-environment matching and ecological filtering for optimal phenotypes on driving community composition (Westoby & Wright, 2006). Second, we assess if the “fingerprints” of climatic changes associated with increased temperatures are revealed in community trait distributions. To make progress on this front, we focus on traits that are proposed to be linked to differences in ecological and evolutionary strategies. Specifically, the leaf-height-seed (LHS) strategy scheme has been proposed to capture the strategy of seed plant species through three trait-related independent dimensions of ecological variation (Westoby, 1998). Variation in vegetative growth, competitive ability, and regenerative processes can be assessed by specific leaf area (SLA; light-capturing area deployed per dry mass allocated), plant canopy height at maturity, and seed mass.

Trait-based ecology has mainly focused on trait-climate linkages by assessing how mean environmental conditions within communities (Kraft et al., 2015) influence the mean community phenotype. This is considered the “mean-field” approach applied at a local scale. Trait-based ecology has largely treated observed community trait variation as reflecting internal niche partitioning driven by species interactions (Keddy, 1992; Weiher & Keddy, 1995; I J Wright, Reich, & Westoby, 2001). However, variation in environmental filtering has also been shown to operate within communities (Adler, Fajardo, Kleinhesselink, & Kraft, 2013). Increasingly, several studies have emphasized that a mean-field approach neglects the importance of intraspecific variation (ITV) within the community (Violle et al., 2012), setting a limit on the generality of resulting ecological inferences. The importance of ITV in ecological studies is undeniable, due to its contribution to functional diversity maintenance (Ross et al., 2017; Siefert et al., 2015) and community assembly processes (Cornwell & Ackerly, 2009; Jung, Violle, Mondy, Hoffmann, & Muller, 2010). Nonetheless, assessing the responses of both intra and interspecific variation could help identify the strength and generality of core assumptions and predictions of trait-based theory for how communities assemble across gradients, and how global warming affects diversity and ecosystem functioning as well.

Recently, Enquist et al. (Enquist et al., 2015, 2017), building on the work of Norberg and colleagues (Norberg et al., 2001), introduced trait driver theory to link how variation in inter- and intraspecific variability in ecological communities can be linked to the performance of individuals across environmental gradients. Trait driver theory (or TDT) goes beyond the mean-field limits (Enquist et al., 2015; Violle et al., 2012) and proposes the investigations of the distribution of the functional trait values by its decomposition into the first four central statistical moments - *mean, variance, skewness, and kurtosis* - to assess the drivers and dynamics of trait distribution in space and time.

### Assessing several prominent assumptions and predictions of trait driver theory (TDT)

According to TDT assumptions, the moments of distributions (including the mean, variance, skewness and kurtosis) are informative about how a species or plant community responds to environmental change. The *mean* value allows us to understand how functional traits shift as the environment alters the optimal trait values. The *variance* provides information about ecological processes affecting trait space wideness. Decreasing variance can be caused by environmental filtering and/or competitive exclusion, whereas competitive niche displacement, immigration and/or temporal variation in the optimum trait value cause the increase of *variance. Kurtosis* measures the “peakedness” of the trait distribution, providing information about community-level ecological processes. TDT predicts that a more peaked and positive distribution reflects competitive exclusion or habitat filtering. Values close to - 1.2 are associated with a uniform trait distribution and even niche partitioning, while more negative values reflect the coexistence of contrasting ecological strategies.

Trait driver theory also makes a series of predictions concerning the dynamical response of communities to environmental change as viewed through the shape of trait distributions. Specifically, in a shifting community, TDT predicts an increase of *skewness* and/or the importance of rare species or rare traits advantage in local coexistence. For example, if phenotype/environment matching is strong, then climate change (e.g., via increases in temperature and/or drought) will cause community trait distributions to track the environment due to decreasing performance or fitness of the existing dominant phenotypes and increasing performance or fitness of some currently rare phenotypes (Enquist et al., 2015, 2017). As this shift occurs, an increase in the skewness is expected, reflecting a lag in time as individuals with newly maladaptive traits slowly decline in frequency. Thus, environmental change (e.g., climate warming/drought) will cause a trait distribution to shift upward or downward along the gradient, and the associated lagging tail will exhibit negative, or positive, skewness relative to the initial distribution, respectively.

By analyzing the trait distribution, one can recast ecological theories that make differing hypotheses about what influences the shape of trait distributions and the ecosystem functioning. These hypotheses include phenotype–environment matching; abiotic filtering; strength of local biotic forces via trait variance and kurtosis; the competitive-ability hierarchy hypothesis; and others (Enquist et al., 2015). In a complementary test for the phenotype-environment matching hypothesis, we also assessed species location in a multidimensional trait space by hypervolume analysis (Blonder, Lamanna, Violle, & Enquist, 2014), which is a feasible way to collapse multi-trait datasets into a simpler framework, allowing us to detect possible shifting of species ecotypes in trait space.

We assessed the predictions of TDT by quantifying species and community level trait distributions across a steep environmental gradient that spans a short geographic distance. This unique system allows us to assess several core assumptions and predictions of TDT. We tested species location and phenotypes shifting in multidimensional hypervolume space along the environmental gradient, to address the following questions: 1) What is the main environmental driver of species distribution? 2) Do plant communities’ trait composition changes along the environmental gradient? 3) Do species phenotypes shift in trait space to match the environmental conditions? 4) How widespread species shift in the multidimensional trait space? 5) What environmental drivers and ecological processes can be unveiled by the shape of the trait distributions? 6) Is there evidence of climate change drivers in *restinga* plant communities?

## Material and Methods

### Study area

Paulo Cesar Vinha State Park (PEPCV) is located between the coordinates 20°33’-20°38’S and 40°23’-40°26’W in Espirito Santo State, Brazil (Figs 1A and B) and holds a prominent position between others *restinga* forests. The PEPCV *restinga* is known for its high environmental heterogeneity, with 11 plant communities spread out side-by-side along narrow environmental gradients (Pereira, 1990). PEPCV is characterized by a conspicuous mosaic of soil types and a slightly waved vertical relief, which creates conditions for the occurrence of lower sites with organic and shallow soils (floodable and permanently flooded areas) and upper sites (non-floodable or drier areas) where well-drained sandy soils predominate.

**Figure 1.**
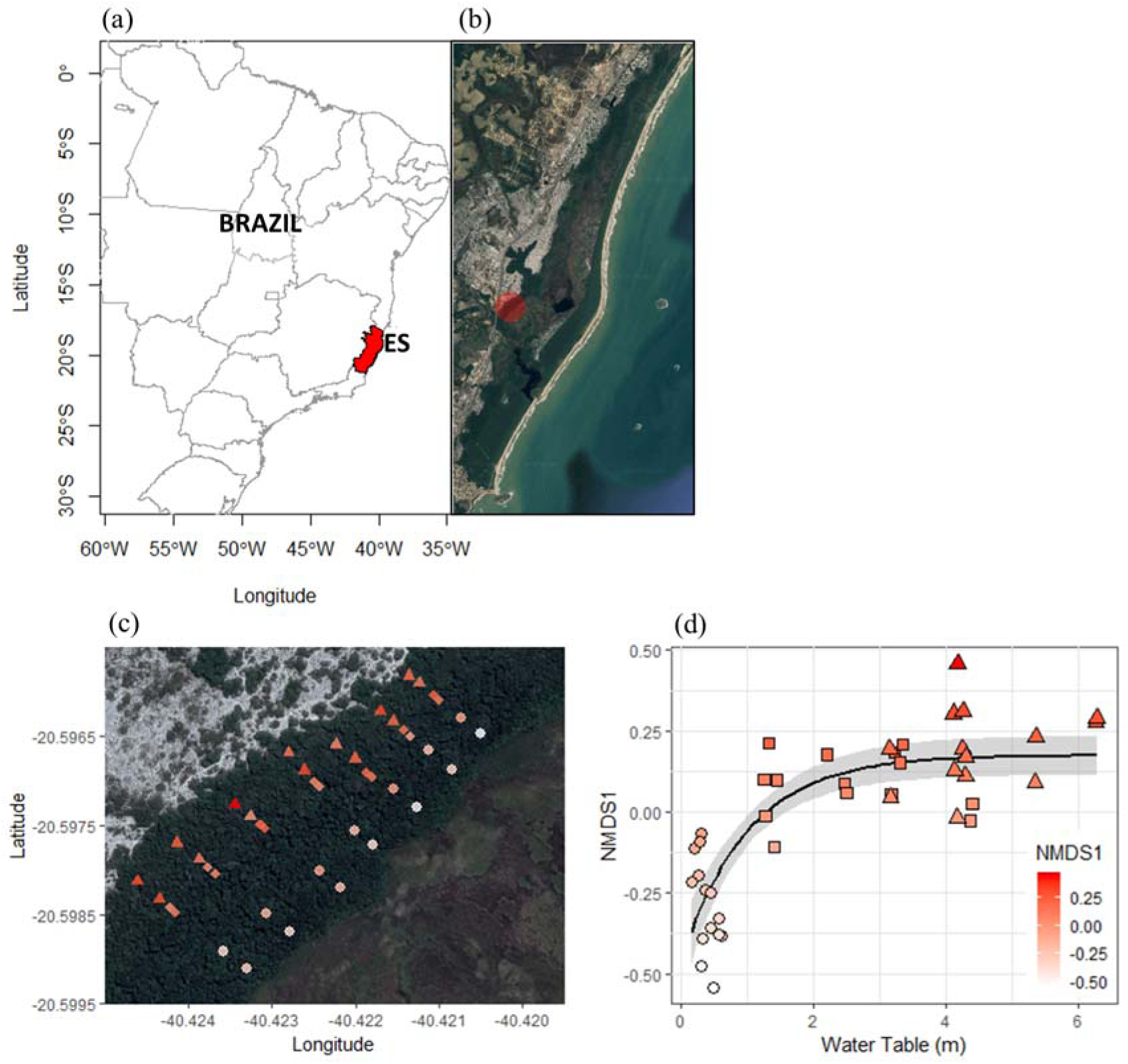
**(a)** Study site consisting of a steep hydrological gradient located in Espirito Santo State (ES) - Brazil. **(b)** Paulo Cesar Vinha State Park and study area location (red dot). (c) Detailed map of the study area, showing the 42 plots/communities that spans a gradient ∼ 207 ± 60 meters from floodable forest (○), intermediate (□) and dry (Δ) forest communities. The variation in symbol color measures the magnitude of community shift in species composition along the NMDS axis (first component eigenvalues) due to the dissimilarity caused by the species replacement, calculated by Simpson index. **(d)** Communities taxonomic composition shifts across the hydrological gradient from the floodable to dry sites, unveiling a steep species replacement along the water table gradient, associated with the shifting of trait composition.

This unique environmental variability makes the Brazilian *restinga* a model system to address timely questions and hypotheses from ecology, such as those related to community assembly processes (Cornwell, Schwilk, & Ackerly, 2006; Keddy, 1992; Stubbs & Wilson, 2004) and the importance of intraspecific variation in a community level context (Bolnick et al., 2011; Jung et al., 2010; Violle et al., 2012; Volf et al., 2016). It could additionally provide an unique opportunity to test the coexistence theories (Hubbell, 2001; Hutchinson, 1957; Macarthur & Levins, 1967) and the effect of the climate change in a wide diversity of environmentally contrasting plant communities, just observed at larger spatial scales (e.g. regional scale).

The study area (Fig. 1C) comprises a short flooding gradient transition, where we set up 42 plots measuring 5 m × 25 m (or 125 m^2^) across three types of forests: floodable, intermediate, and drier forests. All trees with a diameter at breast high (dbh) ≥5cm were tagged, and height and dbh were measured. As the *restinga* plant communities are closely distributed, the fine-scale distribution of the plots allowed us to precisely track the strong effect of the short and continuous soil gradients (distance of 207.4 ± 60.7 meters) in forest taxonomy and trait composition. For simplicity, each plot was defined as a local plant community. The maps were drawn by using the maptools (Bivand & Lewin-Koh, 2017) and raster package (Hijmans, 2017) in the R Software statistical environment (R Core Team, 2018).

### Soil analysis and variable selection

We collected five soil samples per community at a depth of 15 cm, which were then homogenized in the field to produce one compound soil sample for the analysis of the nutritional and physicochemical soil composition from each of 42 local communities. Several parameters were determined by these analyses, including coarseness (proportion of fine and coarse sand, silt, and clay); nutrients (P, K, Na, Ca, Mg, Al, H-Al [potential acidity], Zn, Mn, Cu and B); organic matter (OM); pH; Sodium Saturation Index (SSI), cation exchange capacity (CEC) and base saturation (BS), following the Brazilian Agricultural Research Corporation protocol (Donagema, de Campos, Calderano, Teixeira, & Viana, 2011). We also calculated the soil water retention capacity or field capacity (FC) for each plant community. by collecting two soil samples (at 15 cm depth) in a way that preserves the integrity of the soil structure. We then introduced water to the samples from on end, and installed a fine net in the other to retain the soil. This allowed water flow during the lab experiment, which consisted of carefully introducing water within the pipe until the soil exceeds its maximum water retention capacity. When the water stopped leaking out of the pipe, the samples were weighed in a balance, dried in an oven at 60 □C, until they reached a constant weight. FC was calculated as: FC = (mass of wet soil – mass of dry soil)/mass of wet soil (Donagema et al., 2011).

The water table depth (WT) was directly measured in the floodable areas by digging shallow holes in the soil. Then, we measured the slope variance toward the intermediate and dry plots, taking the WT of the nearby floodable area as a reference. This way, we estimated the WT of the intermediate and the dry plots according to its variance detected along the soil slope. The fitting of the most important environmental variables was performed with generalized additive models (GAM) and generalized linear models (GLM) following a software script provided by Neves et al. (2017), which comprises 1) the exclusion of species found at a single site; 2) the Hellinger transformation of the binary presence/absence data (Legendre & Gallagher, 2001); 3) a forward selection method of environmental variable for redundancy analysis (RDA); and 4) additional and progressive elimination of collinear variables based on their variance inflation factor (VIF) and ecological relevance, until maintaining only those with VIF<4 (Quinn, Keough, & Petraitis, 2002).

Plant species composition matrices were reduced to two dimensions using non-metric multidimensional scaling (NMDS). Ordinations were based on species relative abundance, following the single site species exclusion and Hellinger transformation, as previously described. Communities were compared by the Simpson similarity index, thus, shifting of data points across the NMDS axes is due to the dissimilarity caused by the species replacement. The analysis reached 14.67% of stress. We analyzed the shifting in taxonomy and trait community composition across the WT gradient.

The GAM, GLM, NMDS, and ordination analysis were conducted in the R statistical environment, using the packages vegan (Oksanen et al., 2018), and recluster (Dapporto et al., 2015).

### Trait measurements

The sampling of the leaf and wood traits was performed in over 80% of the most abundant species, according to the species abundance table calculated of the forest inventory we performed within the study area (S.M. Table S2). To maintain the consistency of sampling effort, we kept a similar number of 5 individuals sampled per species, and per environment. Thus, if the species occurs in one, two or three sites, we collected 5, 10 or 15 individuals, to detect the fine-scale environmental influence on species functional traits composition.

We selected plant height (H), specific leaf area (SLA), leaf dry matter content (LDMC) and wood density (WD) because have been highlighted their link to major ecological strategy axes (Chave et al., 2009; Westoby, 1998; Ian J. Wright et al., 2004). Moreover, we also measured leaf thickness (LT) and stomatal density (SD), given their commonly related link with physiological responses to environmental stresses, as water shortage (El-sharkawy, Cock, & Pilar, 1985; Lambers, Chapin, & Pons, 2008). All procedures followed the new handbook protocol for standardized trait measurements (Pérez-Harguindeguy et al., 2013). Leaf trait estimates were based on three leaves per individual, and the same twigs were used to collect the sample for WD measurements. Both leaf and wood samples were previously hydrated and kept in the fridge overnight for 12 hours. We measured leaf area, wet weight, and dry weight to calculate SLA and LDMC. The wood volume was calculated according to the water displacement method (Chave, 2005), with the support of a high precision balance. After that, the twigs were dried in an oven at 60 □C and weighed to determine WD by wood dry mass per wood volume (g/cm^3^).

Stomatal measurements were performed by the leaf imprinting method (Wilson, Pusey, & Otto, 1981), which consist of pushing the leaf surface against a drop of cyanoacrylate adhesive, such a way to leave a thin layer where the leaf surface stay printed. In such a printed layer is possible to account the number and measure the stomatal size or other structures on the leaf surface. In this study, we analyzed both lower and upper leaves surfaces, but the accounting just was performed on the lower surface, as all species exhibited an abaxial pattern of stomata location. We used a microscope with a camera coupled and took photos in 10 times of magnification, which were analyzed in the software TSView, version 6.1.3.2 (Tucsen Imaging Technology Co. Ltd., Fuzhou, Fujian, China).

### Trait predictions

As we measured the height of all individuals reported in the forest inventory, we used that information for the trait prediction procedure. Thus, for each species, we used the trait values collected in the same site of occurrence of the individuals under trait prediction (floodable, intermediate, and dry sites). We took the advantage of the known correlations between the reference trait (e.g. height) and the predicted trait (e.g. SLA). The boundaries of trait variation under prediction were limited to the 95% of confidence interval informed by linear regression. We compared the predicted trait mean-values with the traditional community-weighted mean trait values (Fig. 2), which is calculated by the species traits mean weighted by their abundances in each community. More detailed information and the R script used in trait predictions are provided in the supplementary material (SM), which is available in the online version of this article.

**Figure 2.**
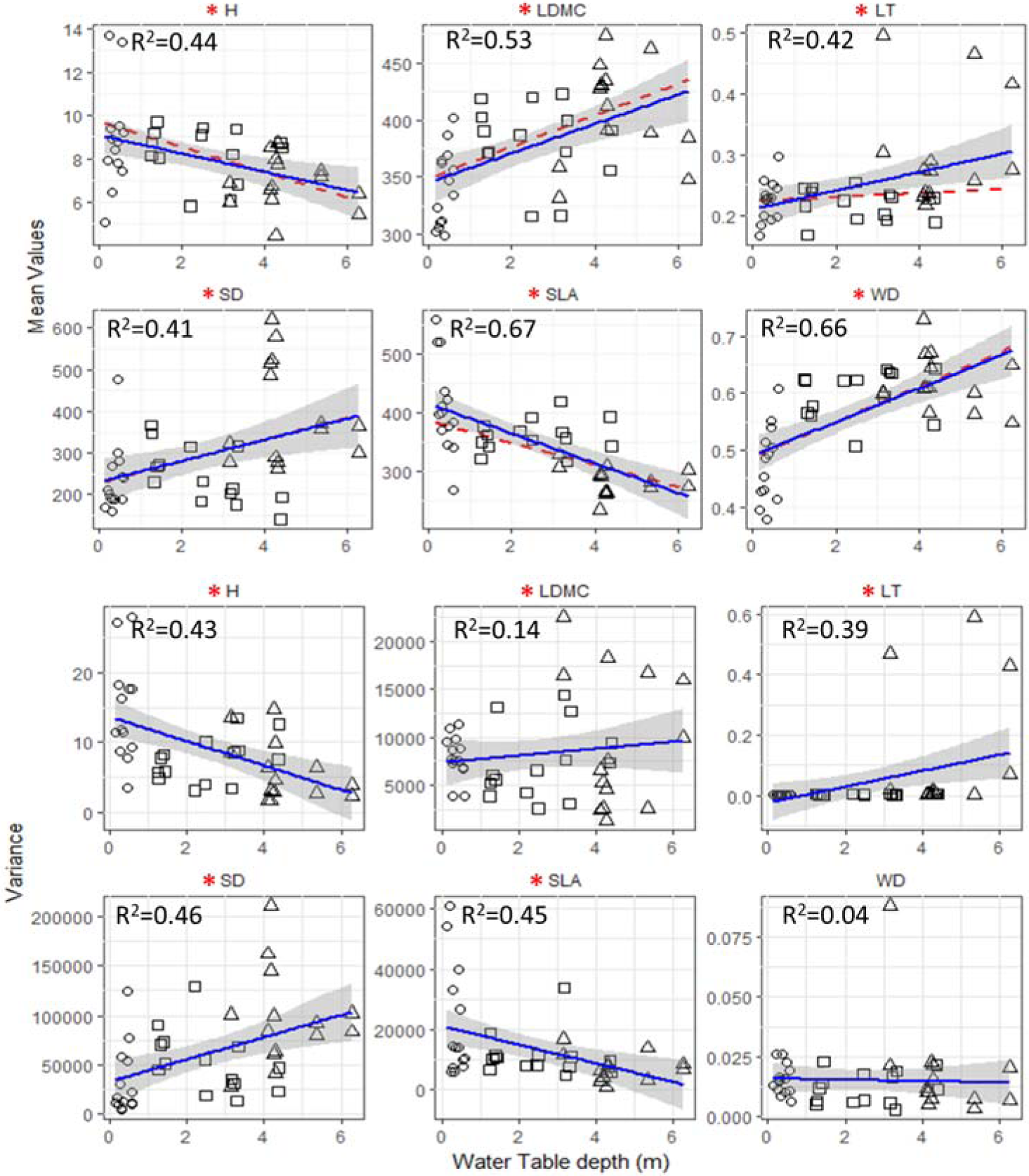
Mean and variance of height, leaf dry matter content (LDMC), leaf thickness (LT), stomatal density (SD), specific leaf area (SLA), and wood density (WD) of 42 plant communities from floodable (○), intermediate (□), and dry (Δ) sites along the water table gradient. The red dashed lines represent the community-weighted means (CWM). Except for LT, the community means calculated by the predicted traits and by weighting species abundance (CWM) fitted closely. Statistically significant at p < 0.05 (*). Despite the short distance, across this steep hydrological gradient, compared to the wet/floodable forests, more dry forests are characterized by shorter trees with dense wood with leaves that are thicker, lower leaf area per unit mass, and higher stomatal density.

### Principal components analysis (PCA) and species hypervolume centroid

The trait dataset (SLA, LDMC, WD, SD, and H values) was collapsed into the PCA space, following the score extraction of the PCA1 and PCA2, which were used to make the hypervolume analysis (Blonder et al., 2014). This procedure is suggested for a multidimensional dataset to avoid issues related to the low number of observations per species, and to optimize the time spent in the hypervolume calculations, as well (Blonder, 2016). We made the hypervolume analysis for each ecotype, that is, for each species in a given site (floodable, intermediate, and dry), and then we extracted the hypervolume centroid, to find the species centroid position into the hypervolume space.

Moreover, the PCA was executed by centering and scaling the score values, as the traits entering the hypervolume analysis must be comparable and uncorrelated (Blonder et al., 2017). The hypervolume analysis was performed in the hypervolume package (Blonder et al., 2014)

## Results

We sampled 863 individuals from 95 species and 73 genera, distributed within 45 families. Myrtaceae (17), Fabaceae (7), Rubiaceae (5) and Sapotaceae (4) were the best-represented families. A large fraction of the trees encountered were identified to genus and species level (96%), and a lower proportion was just identified at the family level (4%; see the species list provided in table S2). Despite the close physical distance between the forest types (207 ±60 meters), we just found six species occurring in all three forest types (*Emmotum nitens, Protium icicariba, Pseudobombax grandiflora, Tapirira guianensis, Aspidosperma pyricollum*, and *Cynophalla flexuosa*).

The forward-selection procedure retained just four environmental variables (WT, B, Mg, coarse sand), explaining 18.05% of the variation in tree species composition, that is, 82% remained unexplained (Table 1). The shifting across the NMDS axes (Figs 1D and E) is given to the dissimilarity caused by the species replacement. Communities were mainly separated in NMDS space by water-related soil parameters (WT and coarse sand).

**Table 1.** Environmental variables selected after the forward selection analysis.

| Variable | adj. $R^2$ cum. | ΔAIC | F | VIF |
| --- | --- | --- | --- | --- |
| Water table depth (WT) | 0.10290 | -12.393 | 5.70 | 1.63 |
| Magnesium (Mg) | 0.14111 | -13.284 | 2.78 | 2.35 |
| Boron (B) | 0.16312 | -13.466 | 2.02 | 2.13 |
| Coarse sand | 0.18057 | -13.471 | 1.81 | 1.83 |

Goodness-of-fit of the predictor variables was assessed through adjusted coefficients of determination, Akaike information criterion (AIC), F-values and significance tests ( $p < 0.01$ in all cases). VIF, variance inflation factor, obtained using the r-squared value of the regression of one variable against all other explanatory variables. adj. $R^2$ cum. = cumulative adjusted coefficient of correlation.

The floodable site has a lower species richness than dry site (S.M.: table S2) and there is a clear species replacement into the first three meters of WT variation (Fig. 1B), that is, communities in the extremities of the gradient are more distinct (Fig. 1A) and flooding exerts a stronger environmental filtering on communities’ taxonomic diversity than do drier conditions. For simplicity, we choose the water table depth (WT) as the main environmental variable because it was the most explicative variable of species distribution and performed the best fitting regression models with all response variables evaluated in this study. All mean trait values shifted from floodable to dry communities, exhibiting a trend of increase in LDMC, SD, WD and LT, and decrease in H and SLA (Fig. 2A). Except for LT, the community-weighted means (CWM) and mean predicted values fitted closely. The drier conditions caused reductions in the variance of H and SLA, whereas LDMC and SD variance increased (Fig. 2B). Few traits showed the trend of changing skewness along the gradient (Fig. 3A). Exceptions for LT, which tended to be more positively skewed; and the communities WD and LDMC values distribution had negative skewness. Both positive (SLA and H) and negative (LDMC and WD) skewness are consistent with the hypothesis of directional shiftings of traits distribution toward drier environmental conditions owned to climate change (Enquist et al., 2017). Moreover, the kurtosis values for the major part of the traits were negative or close to zero (Fig. 3B). Accordingly to TDT, kurtosis values close to - 1.2 may reveal that, in general, the distribution of trait values of all individuals within communities is uniform consistent with niche partitioning. However, we notice a trend of increasing LT toward drier environments. The more dry communities tend to be, the more clustered with more positive kurtosis values for LT than those from floodable communities. The first and second principal components (Fig. 4A) explained 68.5% of trait variation. This variation is mainly due to variation in LDMC (loadings −57.3) and WD (loadings −47.6%) exhibiting the most negative values, and SLA (loadings 55.0) expressing the most positive values. Height (95.2%) was the most positive and determinant loading trait along the PCA2, and stomatal density or SD expressing a negative loading (−27.3%). The traits values distribution of every single individual tree within the PCA space (Fig. 4A) highlights that individuals from the more floodable sites (black symbols) are shifted more in the SLA region of the PCA, whereas those individuals from the drier sites (white symbols) are mainly placed in the quadrant of the trait PCA space influenced more by variation in LDMC, LT, and SD. Individuals are also segregated in PCA space by body size (PC2). The taller ones are mainly from floodable and intermediate sites, whereas smaller individuals are mostly from the dry site.

**Figure 3.**
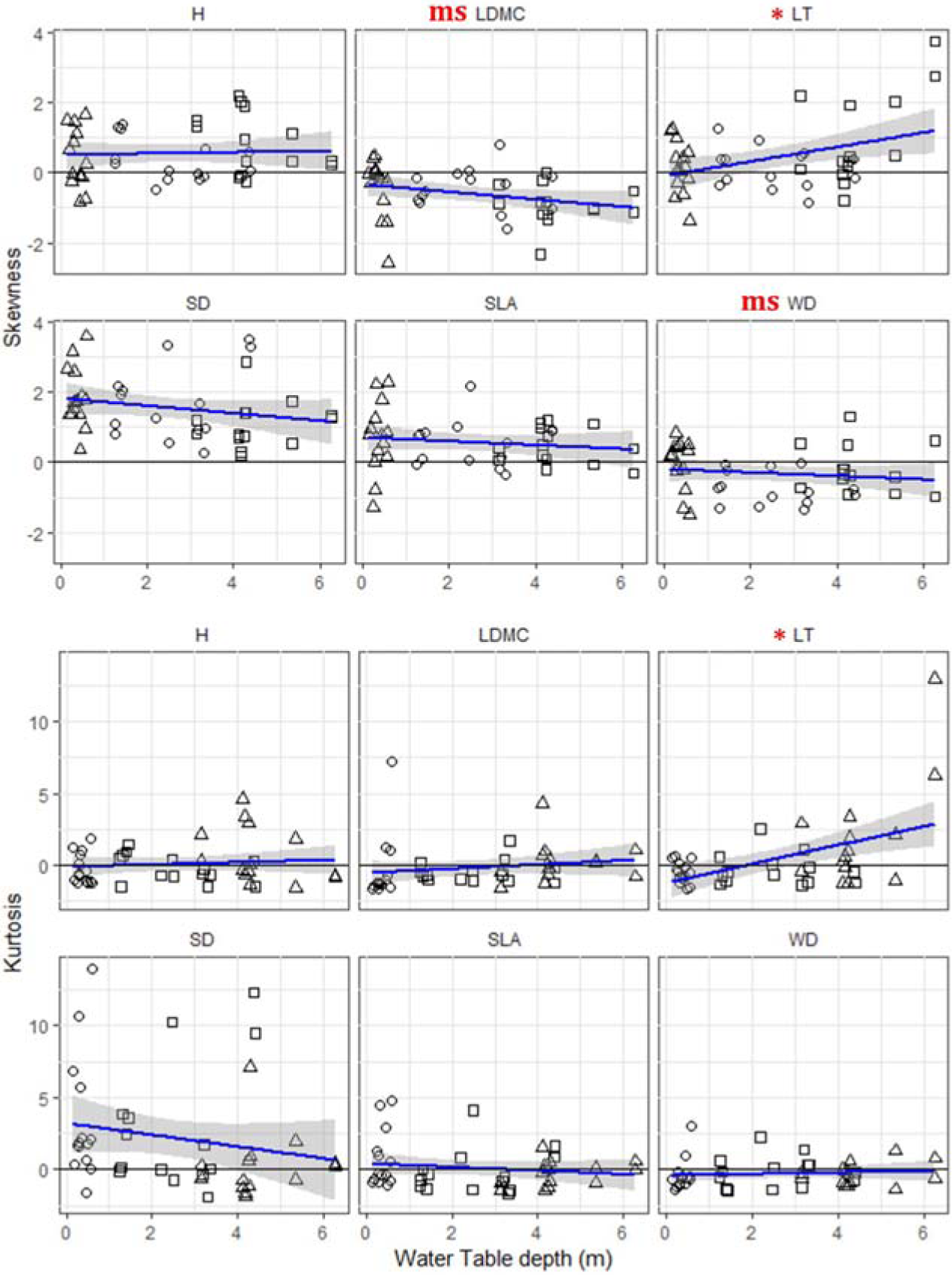
Tests of two assumptions of trait driver theory. Skewness, and kurtosis of height, leaf dry matter content (LDMC), leaf thickness (LT), stomatal density (SD), specific leaf area (SLA), and wood density (WD) of 42 plant communities from floodable (○), intermediate (□), and dry (Δ) sites along the water table gradient. Statistically significant (*) *p* <0.05, or marginally significant (*ms*) [*p*=0.6]. If phenotype/environment matching is strong then environmental change (e.g., climate warming/drought) will cause a trait distribution to shift upward (or downward) across the gradient (Figure 2). The mean trait value of the community will also shift upward (or downward) and the associated lagging tail will exhibit negative (or positive) skewness relative to the initial distribution.

**Figure 4.**
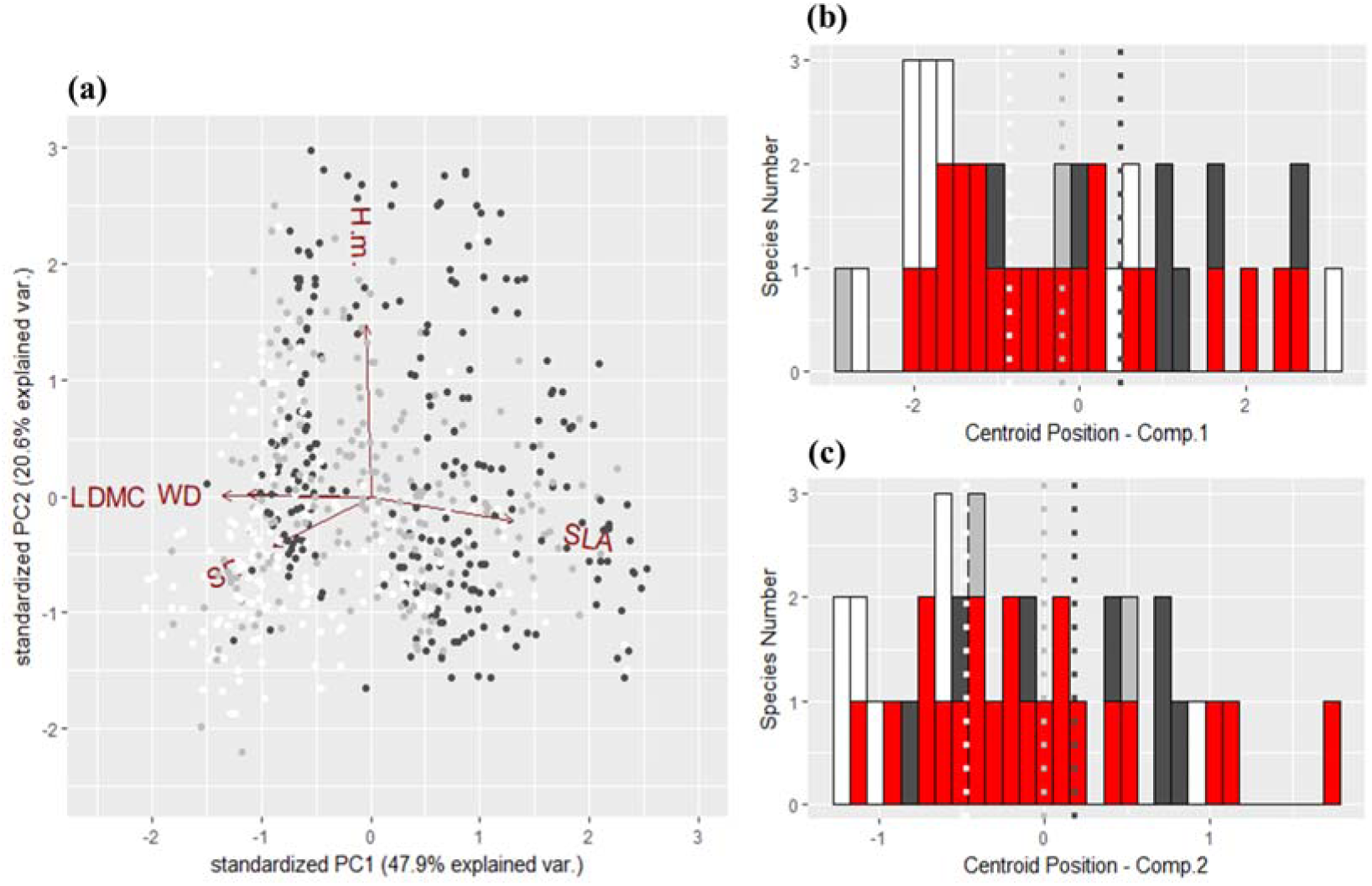
**A** - Principal components analysis (PCA) of specific leaf area (SLA); height (H); wood density (WD); leaf dry matter content (LDMC); and stomatal density (SD) from 626 individual trees within 38 species. **B** and **C** – Positions of species hypervolume centroid, showing specialist species from the floodable (dark gray), intermediate (gray) and dry (white) sites. Generalist species ecotypes (red) have a wider spatial distribution, occurring in two or three sites. Hypervolume centroids were extracted from hypervolume analysis using the PCA scores of the first and second principal components. Dotted lines indicate the mean hypervolume centroids from each site, exhibiting the directional shifting: floodable/intermediate/dry sites (dark gray/gray/white).

Species restricted to one or a few locations along the gradient appear to have well-defined niche space positions. For example, species restricted to either the floodable and dry sites tend to be found in the extremities of trait space, exhibiting positive and negative centroid values, respectively (Figs 4B and C). In contrast, widely-spread species (red bars) exhibit wider distribution along both the centroid axes of the PC1 and PC2, highlighting the role of intraspecific variation in the observed shift of trait composition at the community level (Figs 2A and 5); (Fig. 1D) and the community trait composition received a greater contribution of species ecotypes shifting in trait space. A bidimensional overview of species centroid position in hypervolume space is provided in the box 1, which shows species shifting in a multidimensional hypervolume space by the changing of specific traits or setups of traits.

**Box 1.**
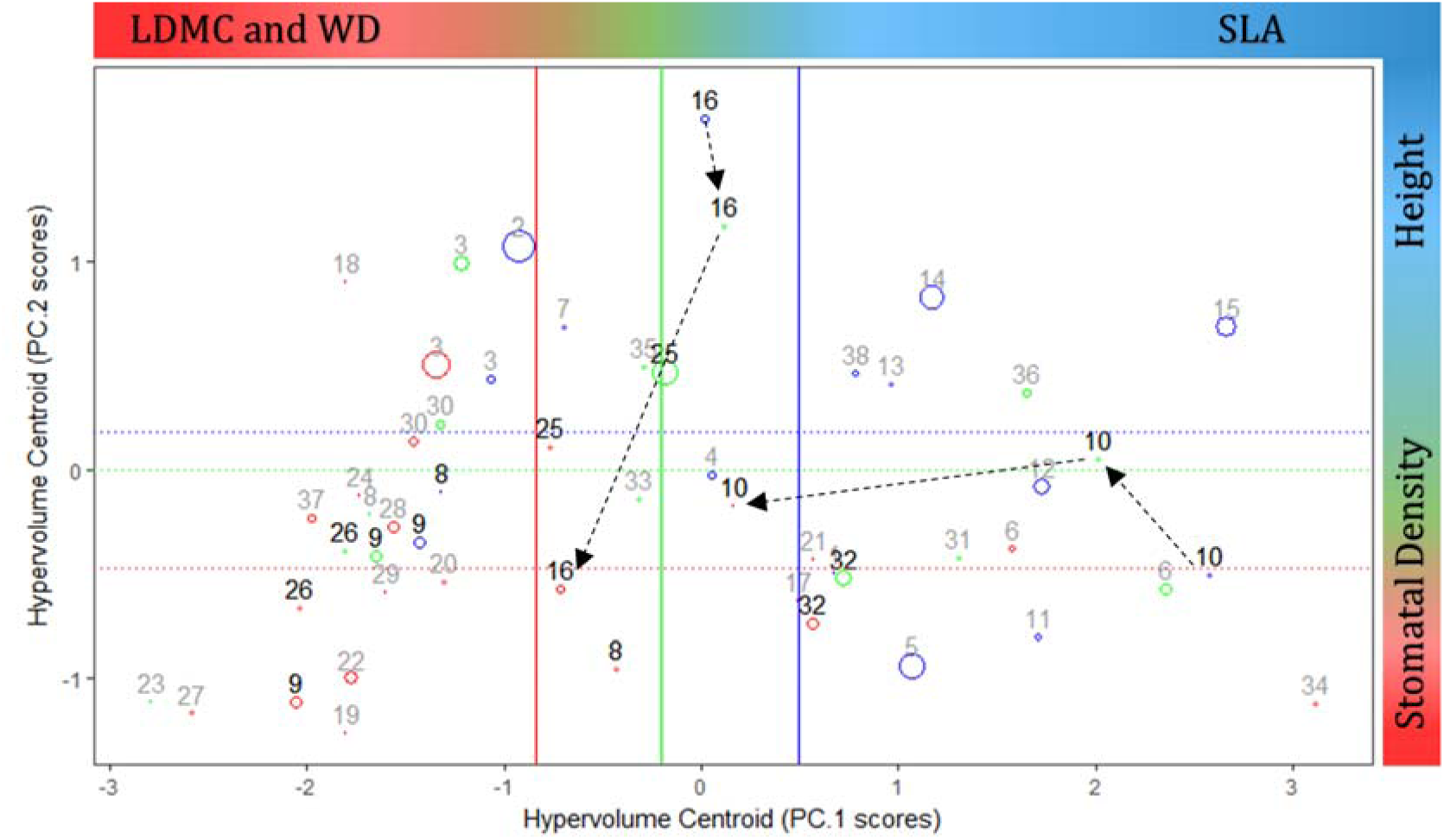
Species distribution along the hypervolume space. Arrows indicate species ecotypes shifting along the two dimensions of the hypervolume, following the shift in the site mean centroid (solid and dotted lines) across floodable (blue), intermediate (green) and dry (red) sites. The first PCA component is related to LDMC, WD, and SLA, whereas the second component is related to H and SD (see Fig. 4A and S.M.: Table S1). The number of individuals within the species is represented by the circle size and the species occurring in more than one site are identified by numbers in bold font. *1 – A. fraxinifolia; 2 – C. brasiliense; 3 – E. nitens; 4 – E. edulis; 5 – G. schotiana; 6 – G. opposite; 7 – I. laurina; 8 – P. heptaphyllum; 9 – P. icicariba; 10 – P. grandiflora; 11 – S. glandulatum; 12 – S. selloi; 13 – S. amara; 14 – S. globulifera; 15 – T. cassinoides; 16 – T. guianensis; 17 – C. arborescens; 18 – B. tetraphylla; 19 – C. guazumifolia; 20 – C. flexuosa; 21 – K. albopunctata; 22 – E. bahiensis; 23 – Eugenia.sp2; 24 – M. cestrifolia; 25 – M. venulosa; 26 – M. bergiana; 27 – M. floribunda; 28 – O. notata; 29 – Ocotea sp; 30 – P. coelomatica; 31 – R. grandiflora; 32 – R. reticulata; 33 – R. capixabensis; 34 – S. sycocapum; 35 – C. saligna; 36 – E. pentaphylla; 37 – Z. glabra; 38 – A. triplinervia*.

## Discussion

The WT was the most important driver of the restinga communities’ taxonomic composition (table 1). The sensitivity to WT was also observed in plant communities from the Amazonian basin (Schietti et al., 2014) and was previously reported the changing of *restinga* forests structure and diversity along flooding gradient (Magnago, Martins, Schaefer, & Neri, 2013). The species replacement (Fig. 1D) was followed by the shifting of the community trait composition. Toward the dry end of the gradient, communities revealed a trend of transition from acquisitive (floodable) to conservative (dry) strategies of resource use (Chave et al., 2009; Ian J. Wright et al., 2004), increasing LDMC, LT, and WD and decreasing SLA and H mean values (Fig. 2A). WD has also been related to soil water availability for its correlation to the wood hydraulic traits (U. Hacke, 2015; U. G. Hacke, Sperry, Pockman, Davis, & McCulloh, 2001), as denser woods with small xylem vessels and high fiber content should be more mechanically resistant to implosion in strong negative water potentials found in drier environments (Swenson & Zambrano, 2017). Similarly, higher stomatal density (SD) seems to be related to the plant water use strategy by a finer control of the leaf transpiration (El-sharkawy et al., 1985). Interestingly, high SD was reported for xeric species (Dunlap & Stettler, 2001; Pearce, Millard, Bray, & Rood, 2005), and our results show the increasing of SD toward the dry communities (Fig. 2), which may contribute to the ongoing discussion about the SD role in saving or luxury water consumption by leaves (Galmés, Flexas, Savé, & Medrano, 2007).

Several environmental drivers have been reported in the literature for constraining trait variance, as elevational gradients (Asner et al., 2017; Sides et al., 2014), high solar radiation, and the shortage of water and soil nutrients (Cornwell & Ackerly, 2009; Katabuchi, Kurokawa, Davies, Tan, & Nakashizuka, 2012). Severe environmental conditions as those found in both extremities of the WT gradient produced filtering effects (Keddy, 1992) on different traits (Fig. 2), reducing the variance in SLA and H (drier plots) and LDMC and SD (wetter plots). Consequently, wet and dry communities have higher diversity of resource acquisition and conservative traits, respectively.

Such shift in trait composition setups with ecological strategies patterns in association to the species replacement (Fig. 2D) was earlier reported by Cornwell and Ackerly (2009) in light and water gradients, and by Katabuchi et al. (2012) in water and soil fertility gradients. The environmental constraints found into the study area seems to exert two majors effects in the *restinga* plant species distribution. First, confining the least plastic species, as the species only located in floodable or drier sites were mainly placed in the extremities of both hypervolume centroid axis (Fig. 4B). Second, widespread species are compelled to vary their setup of traits to fit the changes in the environmental conditions across the strong environmental gradient. Consequently, such species produce phenotypes which fill in a large range of the niche space (hypervolume axes, Fig. 4B, and C), shifting deeply along the multidimensional hypervolume space (Box 1).

These results highlight the importance of analyzing the intraspecific variation in a multi-trait approach, as the species seems to reach out their functional optimality by the variation of specific traits in a multivariate context (Muscarella & Uriarte, 2016). In addition, the leaf and wood traits are not limited to the *economics spectrum* (Laughlin, 2014) and provide a wide variety of ecological strategies by the combination of different traits, which allow tree species to adjust to the environment, reflecting their spatial distribution across environmental gradients. The capacity of a species to produce multiple phenotypes is owed to the phenotypic plasticity (DeWitt, Sih, & Wilson, 1998), which was also reflected in a directional shifting of the shape of species trait distribution, similar to the observed at community level (S.M. Fig. S4).

According to TDT, reflecting selection and ecological filtering for optimal phenotypes, local community trait distributions will tend toward a unimodal distribution and more “peaky” distributions (positive kurtosis), however, more negative distribution with kurtosis close or bellow to −1.2 reflects a uniform distribution, which could reflect the coexistence of contrasting ecological strategies, recent or sudden environmental change (Enquist et al., 2015). Despite some trend of LT trait distribution becoming more “peaky” toward drier communities, our analyses indeed are consistent with such predictions of negative kurtosis being seen predominantly for all traits. The TDT expectation of unimodality of trait distribution was partially fitted, as some communities presented bimodal shapes (S.M. Fig. S4), suggesting the selection of more than one strategy in the restinga communities. On the other hand, the directionality of the shifting in trait composition is consistent with the ecological filtering selection for optimal phenotypes (Enquist et al., 2017).

Similarly to Enquist et al. (2017), the shapes of community trait distribution were characterized by highly positive kurtosis (Fig. 3), indicating strong convergence of functional strategies within a given community with a high number of species or individuals share similar trait values within the communities (Enquist et al., 2015; Gross et al., 2017). Such congested trait space may be the result of strong environmental filtering and/or competitive dominance of individuals with traits closer to the optimum value (Enquist et al., 2017). Interestingly, LT values distribution presented a trend of becoming more peaked toward the dry end of the gradient, which ultimately highlights the importance of species finely adjust such trait in drier environments.

TDT makes two major assumptions about skewed trait distributions, assuming that asymmetries could be caused by 1) the most significative contribution of rare species or rare trait values to the community composition (Enquist et al., 2015); or 2) an ongoing or recent directional shifting in community trait composition due to the changing in an environmental driver (e.g. drought by climate change), resulting in either positive or negative skewed trait distribution depending on the direction that communities shift their mean traits values to track the environmental changing (Enquist et al., 2017). For instance, understory species as *Cyathea pharelatta* depicts the first skewness assumption, as this species contribute with extreme SLA values, which resulted in more positive skewed SLA distribution in floodable communities where this species is present (Fig. S4a).

Our results are consistent with another set of dynamical predictions for how communities respond to climate change. According to TDT if recent directional warming and/or increase in droughts is influencing the species composition of these forests then we would expect to see directional shift in species composition reflected in the trait distributions. In a shifting climate, community trait distributions will shift reflecting a shift in the optimal trait value, but the mean will lag behind the optimal phenotype. In the case where the mean community trait increases as distance to the water table also increase, we would expect that with warming due to climate change, trait distributions will be characterized by *negative* skewness as the community shifts to the new optimal trait value (See scheme in S.M. Fig. S5). In contrast, if the mean community trait is observed to decrease across the water table gradient then warming would lead to communities characterized by *positive* skewness. Thus, the skewness of the trait distribution can reflect shifting optima and past change. In sum, the higher trait moments of communities are expected to deviate in a direction that is dictated by the underlying relationships with access to water.

These forests appear to have experienced unprecedented drought and increased temperatures. Given the close matching between forest functional traits and the hydrological gradient, we would predict that recent directional shifts in climate (associated with water supply and drought) would directionally shift the functional composition of these forests. Indeed, just previous to the field sampling in 2016, this region experienced unprecedented drought and above average temperatures during 2014 and 2015. These were the driest years recorded in the past 50 years (Nobre, Marengo, Seluchi, Cuartas, & Alves, 2016). Further, there has also been a long-term decadal increase/decrease in temperature and drought. ClimateWizard (Girvetz et al., 2009) shows that this area has experienced high rates of change in mean annual temperature over the last 50 years. Moreover, this region is considered a hotspot of climate change in South America (Torres & Marengo, 2014), which is expected to experience severe increases in temperature (above 4°C) and drought events in the upcoming decades (Marengo et al., 2017; Stocker et al., 2013; Torres & Marengo, 2014; van Oldenborgh et al., 2013). Such observed directional shifts in climate associated with water supply would imply that these *restinga* forests may be currently responding.

Several traits do indeed exhibit patterns of relative mean and skewness that are consistent with trait distributions tracking recent climate warming (Fig. 2). This striking result is consistent with the hypothesis that recent climate change is currently causing forests to shift their entire trait distributions in the direction of shifting climates. For example, the observed skewed distributions for SLA and H (two traits whose mean community trait values are negatively associated with depth to the water table) exhibits positive skewness. In contrast, LDMC and WD (two traits that are positively associated with depth to the water table) exhibit negative skewness values. Such signatures in the higher moments of the trait distribution are consistent with the expectation that forests are currently directionally shifting in their trait composition in response to the decreasing of rainfall and the increasing of the temperature, possibly driven by the climate change. Notwithstanding that, a long-term study with periodic taxonomic census and trait collection is needed to test if such skewed patterns are consistent with the shifting of the communities’ trait composition toward drier and warmer environmental conditions. Thus, all plants included in this study were tagged, with the intention of perform a long-term study to track the possible effect of climate-changing in trait community composition along the time.

Local-scale studies in highly diverse environments like *restinga* habitat are timely to accurately assess the microclimate drivers of trait diversity, whose understanding still remains superficial in Ecology (Stark, Lehman, Crawford, Enquist, & Blonder, 2017). Implementing microscale studies to investigate the effect of environmental gradients and climate changing over diversity may provide a more feasible way to track climate changing and answer timely ecological questions regarding to diversity, besides of provide an accurate view of ecological processes and niche-based forces structuring ecological communities. In this study, we used a strong local gradient as evidence that fine-scale processes can drive tree community functional composition and diversity. Our results provide evidence for a strong role of (i) phenotype/environment matching and (ii) functional convergence in shaping the composition of tropical forests and influencing their species distribution and trait composition. Our results also show that the directional shift in species composition is mirrored by a strong shift in ecological strategies reflected by key traits associated with carbon and water use strategies (as also similarly observed by Cornwell and Ackerly (2009) along a topographic gradient of water availability). The moments analysis revealed the possible ecological processes occurring across the WT gradient, which promoted the shifting and filtering of the plant communities’ trait composition, ranging from acquisitive (floodable) to conservative (dry) setup of traits. In addition, the skewness outcome suggests the observed increase in temperature and occurrence of drought is likely already influencing *restinga* plant communities as reflected in observed shift in trait distributions. Our results are consistent with the TDT hypothesis of an ongoing increase in temperature and the drought, matching the climate patterns observed along the last decades in the Southeast region of Brazil.

## Author’s Contributions

JLJ and CRD designed the project; LDT gave support to the taxonomy survey and plant species identification; JLJ analyzed the data; EAN, JLJ and BJE designed the analyses; JLJ, EAN and BJE led the writing of the manuscript. All authors contributed critically to the drafts and gave final approval for publication.

## Acknowledgments

This work was conducted at the University of Arizona and JLJ was financed in part by the Coordenação de Aperfeiçoamento de Pessoal de Nível Superior - Brasil (CAPES) - Finance Code 001 - PDSE program grant n° 88881.131961/2016-01. We are grateful for the field work support of Oberdan Pereira, and the precious advices of Eduardo A. Mattos and Mayara Assis in the initials steps of this study. We are also grateful for the field work assistance of Douglas T. Wandekoken, Felipe Barreto, Fabiano Volponi, Jocimara S. P. Lourenço, Nilton E. Oliveira Filho, Rodrigo Theófilo, José M. L. Gomes, and Danielly hirata. The authors declare no conflicts of interest.

## Supplementary Material

Additional Supporting Information may be found in the online version of this article:

**Appendix S1.** Trait prediction

**Appendix S2.** PCA loadings and species shifting in two-dimensional centroid space

**Appendix S3.** Shapes of the traits distribution

**Coding.** Pred_trait.R function and text file script for the trait predictions

